# Neutralization of SARS-CoV-2 EG.5/EG.5.1 by sera from ZF2001 RBD-dimer and its next-generation vaccines

**DOI:** 10.1101/2023.09.02.556038

**Authors:** Yaling An, Xuemei Zhou, Lifeng Tao, Haitang Xie, Chenxi Yang, Dedong Li, Ruyue Wang, Hua Hu, Kefang Liu, Lianpan Dai, Kun Xu, George F. Gao

**Affiliations:** Savaid Medical School, University of Chinese Academy of Sciences, Beijing, China; CAS Key Laboratory of Pathogen Microbiology and Immunology, Institute of Microbiology, Chinese Academy of Sciences, Beijing, China; School of Life Sciences, Hebei University, Baoding, Hebei, China; Anhui Zhifei Longcom Biopharmaceutical Co. Ltd, Hefei, Anhui, China; Yijishan Hospital of Wannan Medical College, Wuhu, Anhui, China; College of Veterinary Medicine, China Agricultural University, Beijing, China; Research Network of Immunity and Health (RNIH), Beijing Institutes of Life Science, Chinese Academy of Sciences, Beijing, China

## Abstract

SARS-CoV-2 Omicron EG.5 and EG.5.1 are surging in several areas of the world, including China. Compared with XBB.1, EG.5 contains additional mutations of F456L and S486P in the spike protein receptor binding domain (RBD) and its subvariant EG.5.1 carries a further spike mutation Q52H. The immune escape potential of EG.5/EG.5.1 is of great concern. In this study, we evaluated the neutralization activities of sera from participants who received COVID-19 inactivated vaccines, protein subunit vaccine ZF2001 or a booster vaccination of Delta-BA.5 RBD-heterodimer protein vaccine, and participants who had a breakthrough infection during a wave of BF.7/BA.5.2 circulation in December 2022. Neutralization profiles elicited by bivalent RBD-heterodimer vaccine candidates containing XBB.1.5 antigen were evaluated in a murine model. We found that EG.5 and EG.5.1 displayed similar immune evasion potential to XBB.1 and XBB.1.5. The Delta-BA.5 RBD-heterodimer booster induced higher neutralizing titers against the tested XBB subvariants, including EG.5 and EG.5.1, than breakthrough infection by BF.7 or BA.5.2. In addition, Delta-XBB.1.5 and BQ.1.1-XBB.1.5 RBD-heterodimer vaccines induced high neutralizing activities against XBB sub-variants in a murine model, suggesting that next-generation COVID-19 vaccines with updated components must be developed to enhance the protection efficacy against the circulating SARS-CoV-2 strains.

## Main text

Severe acute respiratory syndrome coronavirus 2 (SARS-CoV-2) variants are continuously evolving, emerging and circulating. Recently, XBB.1.9.2.5 (EG.5) and XBB.1.9.2.5.1 (EG.5.1) are surging in several areas of the world^1^. EG.5/EG.5.1 have become the most prevalent SARS-CoV-2 variants in China since August 2023, accounting for more than 70% of the virus sequences. The World Health Organization (WHO) risk evaluation determined a moderate growth advantage of EG.5, and designated it as the variant of interest (VOI) on 9 Aug 2023, accompanied by the other two VOC (XBB.1.5 and XBB.1.16)^1^. Compared with XBB.1, EG.5 contains additional mutations of F456L and S486P in the spike (S) protein receptor binding domain (RBD). Its subvariant EG.5.1 carries a further spike mutation Q52H on the N-terminal domain of S protein (Fig. 1a; Supplementary information, Fig. S1). Therefore, the immune escape potential of EG.5/EG.5.1 is of great concern.

**Fig. 1:**
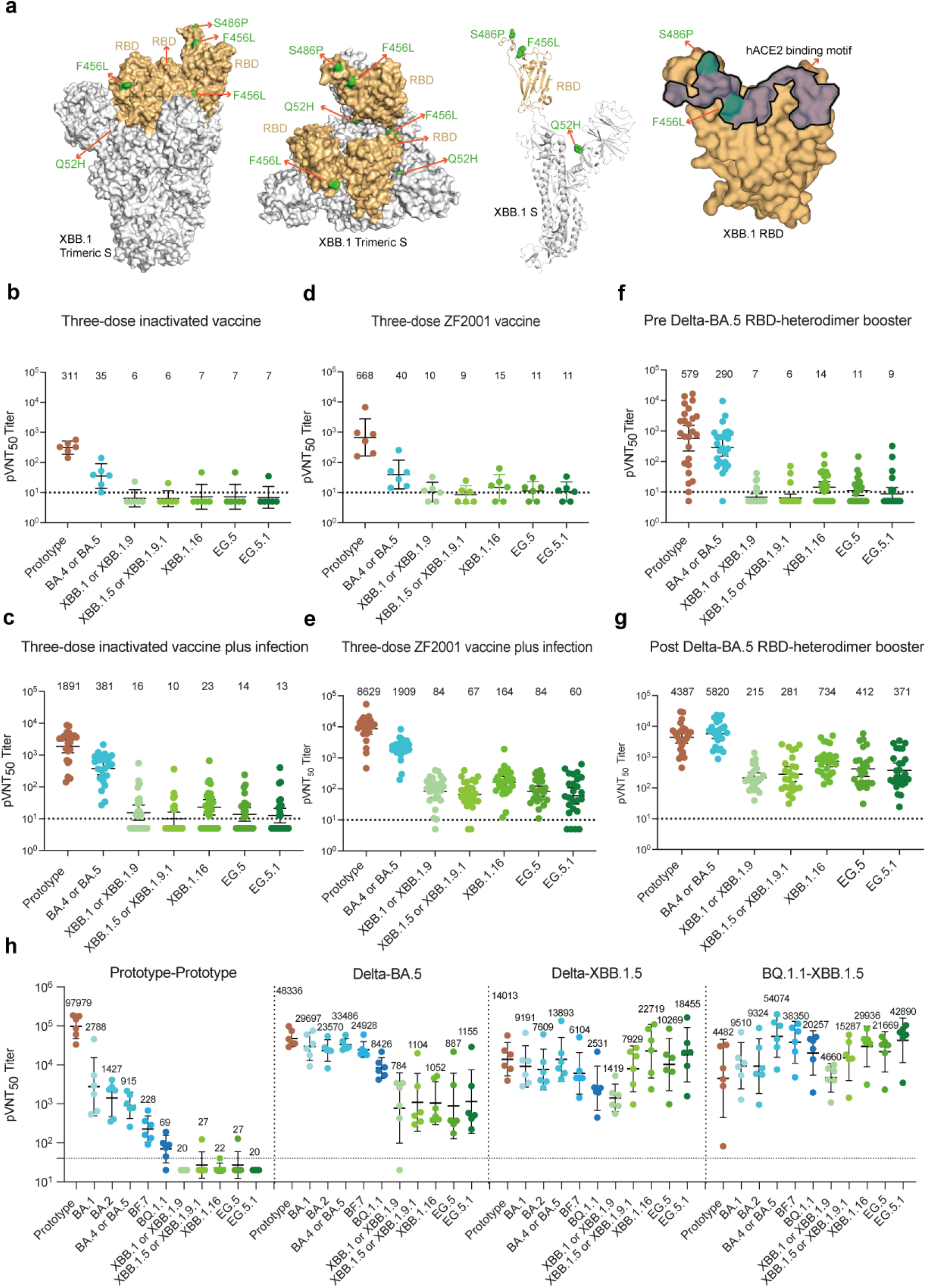
Neutralizing antibodies against SARS-CoV-2 variants by sera from humans and mice. (a)Additional mutations F456L, S486P and Q52H in EG.5.1 S protein compared to XBB.1. XBB.1 S protein is shown as gray color (PDB: 8IOU^9^). RBD is shown as gold color. Footprint of hACE2 on SARS-CoV-2 RBD is highlighted in brown. Residues that mutated in Omicron EG.5.1 subvariant are colored in green.(b)The pVNT_50_ in participants who had received three doses of inactivated vaccine. (c)The pVNT_50_ in participants who had breakthrough infections during the late-2022 omicron subvariant BF.7 and BA.5.2 wave after three doses of inactivated vaccine. (d)The pVNT_50_ in participants who had received three doses of ZF2001 vaccine. (e)The pVNT_50_ in participants who had breakthrough infections during the late-2022 omicron subvariant BF.7 and BA.5.2 wave after three doses of ZF2001 vaccine. (f-g) The pVNT_50_ in participants who had received a booster of Delta-Omicron BA.5 RBD-heterodimer vaccine. The sera were collected on the same day pre-boosting vaccination (f) and about three weeks post-boosting vaccination (g). (h) Groups of 6-to 8-week-old female BALB/c mice (n = 6) were immunized with three doses of prototype-prototype, Delta-BA.5, Delta-XBB.1.5 or BQ.1.1-XBB.1.5 RBD-dimer vaccine using AddaVax as adjuvant, 21 days apart. PBS plus adjuvant was given as the sham control. Sera were collected at 14 days post the third immunization for pVNT_50_ titration.

SARS-CoV-2 herd immunity has been established through coronavirus disease 2019 (COVID-19) vaccine inoculation^2^. Inactivated vaccines (CoronaVac and BBIBP-CorV) are widely used in China and many other countries^3^. The protein subunit vaccine ZF2001, based on the immunogen of SARS-CoV-2 RBD-homodimer, has been authorized in China, Uzbekistan, Indonesia, Colombia, Kenya and Belarus^4, 5^. However, breakthrough infections occurred due to the waning immunity, immune evasion by the SARS-CoV-2 variants, and high transmissibility of Omicron subvariants. The immune responses induced by vaccination and breakthrough infection against EG.5/EG.5.1 need to be elucidated. In addition, we previously designed the immunogen of SARS-CoV-2 RBD-heterodimers as next-generation vaccines of ZF2001 to elicit broad immunity against variants^6^. Recently, Delta-Omicron BA.5 RBD-heterodimer vaccine (ZF2202-A) has been under the clinical trial study (NCT05850507). Here, we evaluated the neutralizing activities of Delta-BA.5 vaccine elicited human sera and the other next-generation vaccine candidates, such as Delta-XBB.1.5 and BQ.1.1-XBB.1.5, elicited murine sera. Therefore, the samples included in this study were from (1) participants who received three homologous doses of COVID-19 inactivated vaccines or ZF2001 vaccine; (2) participants with breakthrough infection during a wave of BF.7/BA.5.2 circulation in Beijing in December 2022; (3) participants who received a booster vaccination of Delta-BA.5 RBD-dimer protein vaccine; (4) mice immunized with three doses of bivalent RBD-heterodimer vaccines. The detailed information of the participants is available in Supplementary information, Table S1. We evaluated the neutralizing activities using a penal of vesicular stomatitis virus backbone-based pseudotyped viruses displaying SARS-CoV-2 S protein.

Three homologous doses of COVID-19 inactivated vaccines or ZF2001 vaccine-induced neutralizing GMTs of 311 and 668 against prototype pseudovirus, respectively (Fig. 1b, d). However, XBB.1.5, XBB.1.16, EG.5 and EG.5.1 showed significant resistance to the vaccine-elicited neutralizing immunity, with geometric mean titers (GMTs) near or below the limit of detection (LOD) (Fig. 1b, d; Supplementary information, Fig. S2). Breakthrough infection after three doses of inactivated vaccines immunization substantially enhanced the neutralizing responses (Fig. 1c). By contrast, three doses of ZF2001 vaccination plus breakthrough infection induced higher neutralizing GMTs (BA.5: 1909; XBB.1: 84; XBB.1.5: 67; XBB.1.16: 164; EG.5: 84; EG.5.1: 60) and seropositive rates (BA.5: 100%; XBB.1: 96%; XBB.1.5: 92%; XBB.1.16: 100%; EG.5:100%; EG.5.1: 84%) (Fig. 1e; Supplementary information, Fig. S3).

The study on bivalent Delta-BA.5 RBD-dimer vaccine revealed a large enhancement in neutralizing activities against the pseudoviruses by a factor of 7.6–52.4 post-boosting (Fig. 1f, g). A booster dose of Delta-BA.5 RBD-dimer vaccine induced 100% neutralization seropositive rates against all the tested pseudoviruses with GMTs of 4387, 5820, 215, 281, 734, 412 and 371 against prototype, BA.5, XBB.1, XBB.1.5, XBB.1.16, EG.5 and EG.5.1, respectively (Fig.1g; Supplementary information, Fig. S3). In addition, the neutralizing GMTs against BA.5, XBB1.5/XBB.1.16 and EG.5/EG.5.1 induced by Delta-BA.5 RBD-dimer vaccine booster were significantly higher than the breakthrough infection after three doses of inactivated vaccine or ZF2001 (Fig.1 c, e, g; Supplementary information, Fig. S4). These results indicate that the bivalent Delta-BA.5 RBD-heterodimer vaccine elicits broad neutralizing activities in humans, making it a promising booster vaccine candidate.

Next, we analyzed the neutralization profiles induced by several RBD-heterodimer vaccines in a murine model. The prototype-prototype RBD-dimer elicited a neutralizing titer as high as approximately 10^5^ against the prototype pseudovirus. Omicron subvariants BA.1, BA.2, BA.5, BF.7 and BQ.1.1 escape the neutralizing responses with gradually increased severity. The neutralization activities against all the XBB subvariants, including EG.5/EG.5.1, decreased to background levels (Fig. 1h). Compared to the prototype-prototype RBD-dimer, Delta-BA.5 induced a broader neutralization profile, with neutralizing GMTs that ranging between 23570 and 48336 against prototypes and Omicron BA.1/BA.2/BA.5/BF.7, 8426 against BQ.1.1, and between 784 and 1155 against XBB.1/XBB.1.5/XBB.1.16 and EG.5/EG.5.1 (Fig. 1h). Moreover, Delta-XBB.1.5 and BQ.1.1-XBB.1.5 RBD-dimer vaccines induced higher neutralizing activities against XBB sub-variants than Delta-BA.5, suggesting that using an XBB.1.5-containing RBD heterodimer is beneficial to prevent XBB sub-variants, including EG.5 and EG.5.1 (Fig. 1h).

These data revealed that the XBB.1.5, XBB.1.16, EG.5 and EG.5.1 escaped the antibody responses induced by prototype SARS-CoV-2-based vaccines with similar severity. Omicron subvariants BF.7/BA.5.2 breakthrough infection upon prior three-dose of ZF2001 vaccination elicited substantially neutralizing activities against XBB sub-variants, including EG.5 and EG.5.1, with more than 84% seropositive rates (Fig. 1e; Supplementary information, Fig. S3). By comparison, a booster of the Zhifei’s third-generation COVID-19 vaccine Delta-BA.5 RBD-heterodimer induced higher neutralizing titers against EG.5 and EG.5.1 in humans, demonstrating the need to update the vaccine immunogen (Fig. 1g). Furthermore, the other COVID-19 next-generation vaccine candidates, Delta-XBB.1.5 and XBB-BQ.1.1 RBD-heterodimers, were evaluated in a murine model and elicited broad-spectrum neutralizing activities against SARS-CoV-2 variants, particularly the currently-circulating EG.5 and EG.5.1, which should be further clinically developed (Fig. 1h).

In order to prevent the diseases caused by SARS-CoV-2 infection, boosting vaccination is crucial, especially for vulnerable populations such as the elderly, immunocompromised individuals and people suffering chronic diseases like diabetes and hypertension. Moreover, the risks of sequelae in multiple organ systems caused by SARS-CoV-2 infection can persist for two years^7^. The cumulative burden of health loss of people infected by SARS-CoV-2 should be concerned. The SARS-CoV-2 vaccines protect against COVID-19, reduce the symptoms caused by breakthrough infection, and reduce the risk of long-term health effects following infection^8^. Therefore, next-generation COVID-19 vaccines with updated components must be developed to enhance the protection efficacy against the circulating SARS-CoV-2 strains and long COVID-19.

The pVNT_50_ is shown as the geometric mean titer at the top in each panel. The error bars indicate 95% confidence intervals. The dashed horizontal line indicates the limit of detection (LOD). A pVNT_50_ value below the LOD was determined as half the LOD.

