## Supplemental information for "Neutralization of SARS-CoV-2 EG.5/EG.5.1 by sera from ZF2001 RBD-dimer and its next-generation vaccines"

### Supplementary information

### Supplementary information

### Materials and Methods

#### Human serum samples

Human serum samples were classified into fifth groups in this study. The first group received three doses of inactivated vaccine (CoronaVac or BBIBP-CorV). The second group received three doses of inactivated vaccine and had breakthrough infection during the BF.7/BA.5.2 wave in late 2022, Beijing, China. The third group received three doses of recombinant protein subunit vaccine (ZF2001). The fourth group received three doses of ZF2001 vaccine and had breakthrough infection during the BF.7/BA.5.2 wave in late 2022, Beijing, China. The fifth group received two or three doses of inactivated vaccine, followed by boosting with one dose of Delta-Omicron BA.5 RBD-heterodimer protein vaccine (ZF2202-A). For the fifth group, blood samples were collected on the same day before boosting immunization and about three weeks after receiving the booster vaccine. Serum samples from the fifth group were from a clinical trial (NCT05850507), and were approved by the clinical research ethics board of the First Affiliated Hospital of Wannan Medical College (Yijishan Hospital) ([2023]KY34). Samples from the first group were provided by Anhui Zhifei Longcom Biopharmaceutical. Samples from the second, third and fourth groups were collected from the participants in real-world. All participants signed the written informed consent. Detailed information is available in Table S1.

#### Pseudotyped virus neutralization assay

The pseudotyped virus displaying SARS-CoV-2 (prototype and variants) S protein expresses GFP in infected cells. They were prepared as previously described[1]. Serially diluted mice sera were incubated with pseudotyped virus at 37°C for 1 hour, then the sera-virus mixtures were transferred to pre-plated Vero

E6 cell monolayers in 96-well plates. After incubation for 15 hours, the transducing unit numbers were calculated on a CQ1 confocal image cytometer (Yokogawa). Neutralization titer was determined by fitting nonlinear regression curves using GraphPad Prism and calculating the reciprocal of the serum dilution required for 50% neutralization of infection. Neutralization titer below the limit of detection was determined as half the limit of detection.

**Mouse experiments**

Specific pathogen-free (SPF) female BALB/c mice were purchased from Beijing Vital River Laboratory Animal Technology Co., Ltd. (licensed by Charles River). All mice were allowed free access to water and standard chow diet and provided with a 12-hour light and dark cycle (temperature: 20-25°C, humidity: 40%-70%). All mice used in this study are in good health and are not involved in other experimental procedure. They were housed under SPF conditions in the laboratory animal facilities at IMCAS. The mice experiments conducted in IMCAS were approved by the Committee on the Ethics of Animal Experiments of the IMCAS, and performed in compliance with the recommendations in the Guide for the Care and Use of Laboratory Animals of the IMCAS Ethics Committee.

Antigen proteins were prepared as previously described[2]. AddaVax adjuvant was purchased from InvivoGen. Groups (n = 6) of 6- to 8-week-old female BALB/c mice were immunized with three doses of 2-μg immunogens adjuvanted by AddaVax. The interval between the first and second doses was 21 days. The interval between the second and third doses was 21 days. The serum samples were collected 14 days after the last immunization.

**Statistical analysis**

Pseudovirus neutralization titer ($pVNT_{50}$) was determined by fitting nonlinear regression curves using GraphPad Prism and calculating the reciprocal of the serum dilution required for 50% neutralization of infection. The values shown are GMT ±95% confidence interval (CI). p values shown in Supplementary information, Figs. S2 and S4 were analyzed with a Kruskal-Wallis test first, and then Dunn's multiple comparison test when the Kruskal-Wallis test was rejected.

### Table S1. Demographic and characteristics of participants

| Groups | Group 1 | Group 2 | Group 3 | Group 4 | Group 5 | |
|---|---|---|---|---|---|---|
| **Status of vaccination or SARS-CoV-2 infections** | Three-dose inactivated vaccine | Three-dose inactivated vaccine plus breakthrough infection* | Three-dose ZF2001 | Three-dose ZF2001 plus breakthrough infection | Pre Delta-BA.5 RBD-heterodimer booster | Post Delta-BA.5 RBD-heterodimer booster |
| **No. of participants** | 6 | 25 | 6 | 25 | 25 | |
| **Age, Years** | | | | | | |
| **Mean (SD)**<br>**Median (IQR)** | 29.7 (7.8)<br>27.0 (24.8-33.8) | 29.7 (8.5)<br>27.0 (24.0-30.0) | 34.2(10.4)<br>30.5 (26.5-42.5) | 35.8 4.2(14.4)<br>31.0 (25.5-37.5) | 48.3(17.6),<br>54.0 (32-65.0) | |
| **Male, n (%)** | 4.0（66.7%） | 6.0 (24.0%) | 2.0（33.3%） | 11.0 (44.0%) | 15.0 (60.0%) | |
| **Female, n (%)** | 2.0（33.3%） | 19.0 (76.0%) | 4.0（66.7%） | 14.0 (56.0%) | 10.0 (40.0%) | |
| **Time interval between last dose and breakthrough infection, Days** | | | | | | |
| **Mean (SD)**<br>**Median (IQR)** | N.D.** | 359.5 (90.2)<br>396.0 (361.0-406.0) | N.D. | 485.6 (141.7)<br>484.0 (394.5,561.0) | N.D. | |
| **Time interval between the last event (last dose or breakthrough infection) and blood sampling, Days** | | | | | | |
| **Mean (SD)**<br>**Median (IQR)** | 15.7 (1.0)<br>16.0 (14.8,16.2) | 33.5 (9.3)<br>32.0 (28.0,35.5) | 27.8 (30.9)<br>14.0 (14.0-48.5) | 25.2 (4.4)<br>25.0 (22.0,28.5) | 479.4.8 (138.6)<br>534.0 (372.5-557.5) | 22.6 (1.6)<br>23.0 (21.0-24.0) |

*Breakthrough infection were confirmed by antigen test for their nasopharyngeal swabs

**N.D.: not determined

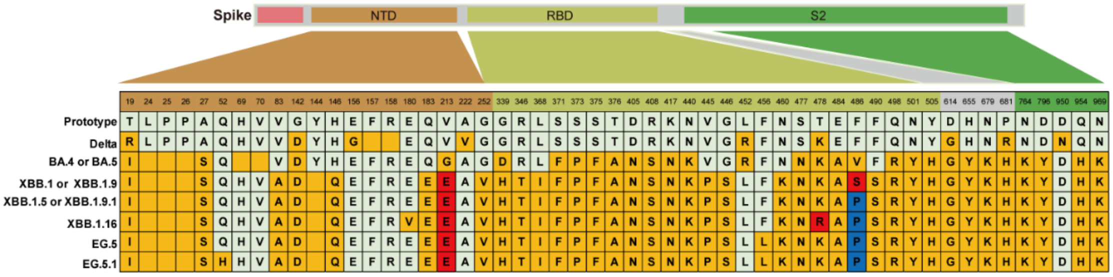

**Figure S1. Mutations in the spike protein of prototype SARS-CoV-2 and variants.**

The amino acid differences in SARS-CoV-2 variant sequences from those in the prototype isolate, including substitutions and deletions, are highlighted in gold. The second and third type of amino acid on the same locus is marked in red and blue. Deletion mutations are shown as blank squares.

| Three-dose inactivated vaccine | | | | | | | |
|---|---|---|---|---|---|---|---|
| | Prototype | BA.4 or BA.5 | XBB.1 or XBB.1.9 | XBB.1.5 or XBB.1.9.1 | XBB.1.16 | EG.5 | EG.5.1 |
| Prototype | | | | | | | |
| BA.4 or BA.5 | >0.9999 | | | | | | |
| XBB.1 or XBB.1.9 | 0.0032 | 0.2892 | | | | | |
| XBB.1.5 or XBB.1.9.1 | 0.0029 | 0.2685 | >0.9999 | | | | |
| XBB.1.16 | 0.0054 | 0.4151 | >0.9999 | >0.9999 | | | |
| EG.5 | 0.0049 | 0.3866 | >0.9999 | >0.9999 | >0.9999 | | |
| EG.5.1 | 0.0036 | 0.3113 | >0.9999 | >0.9999 | >0.9999 | >0.9999 | |

| Three-dose inactivated vaccine plus infection | | | | | | | |
|---|---|---|---|---|---|---|---|
| | Prototype | BA.4 or BA.5 | XBB.1 or XBB.1.9 | XBB.1.5 or XBB.1.9.1 | XBB.1.16 | EG.5 | EG.5.1 |
| Prototype | | | | | | | |
| BA.4 or BA.5 | >0.9999 | | | | | | |
| XBB.1 or XBB.1.9 | <0.0001 | <0.0001 | | | | | |
| XBB.1.5 or XBB.1.9.1 | <0.0001 | <0.0001 | >0.9999 | | | | |
| XBB.1.16 | <0.0001 | 0.0006 | >0.9999 | >0.9999 | | | |
| EG.5 | <0.0001 | <0.0001 | >0.9999 | >0.9999 | >0.9999 | | |
| EG.5.1 | <0.0001 | <0.0001 | >0.9999 | >0.9999 | >0.9999 | >0.9999 | |

| Three-dose ZF2001 vaccine | | | | | | | |
|---|---|---|---|---|---|---|---|
| | Prototype | BA.4 or BA.5 | XBB.1 or XBB.1.9 | XBB.1.5 or XBB.1.9.1 | XBB.1.16 | EG.5 | EG.5.1 |
| Prototype | | | | | | | |
| BA.4 or BA.5 | >0.9999 | | | | | | |
| XBB.1 or XBB.1.9 | 0.0121 | 0.6845 | | | | | |
| XBB.1.5 or XBB.1.9.1 | 0.0034 | 0.2842 | >0.9999 | | | | |
| XBB.1.16 | 0.1763 | >0.9999 | >0.9999 | >0.9999 | | | |
| EG.5 | 0.0334 | >0.9999 | >0.9999 | >0.9999 | >0.9999 | | |
| EG.5.1 | 0.0203 | 0.9679 | >0.9999 | >0.9999 | >0.9999 | >0.9999 | |

| Three-dose ZF2001 vaccine plus infection | | | | | | | |
|---|---|---|---|---|---|---|---|
| | Prototype | BA.4 or BA.5 | XBB.1 or XBB.1.9 | XBB.1.5 or XBB.1.9.1 | XBB.1.16 | EG.5 | EG.5.1 |
| Prototype | | | | | | | |
| BA.4 or BA.5 | >0.9999 | | | | | | |
| XBB.1 or XBB.1.9 | <0.0001 | <0.0001 | | | | | |
| XBB.1.5 or XBB.1.9.1 | <0.0001 | <0.0001 | >0.9999 | | | | |
| XBB.1.16 | <0.0001 | 0.0041 | >0.9999 | 0.6273 | | | |
| EG.5 | <0.0001 | <0.0001 | >0.9999 | >0.9999 | >0.9999 | | |
| EG.5.1 | <0.0001 | <0.0001 | >0.9999 | >0.9999 | 0.8299 | >0.9999 | |

| Pre Delta-BA.5 RBD-heterodimer booster | | | | | | | |
|---|---|---|---|---|---|---|---|
| | Prototype | BA.4 or BA.5 | XBB.1 or XBB.1.9 | XBB.1.5 or XBB.1.9.1 | XBB.1.16 | EG.5 | EG.5.1 |
| Prototype | | | | | | | |
| BA.4 or BA.5 | >0.9999 | | | | | | |
| XBB.1 or XBB.1.9 | <0.0001 | <0.0001 | | | | | |
| XBB.1.5 or XBB.1.9.1 | <0.0001 | <0.0001 | >0.9999 | | | | |
| XBB.1.16 | <0.0001 | 0.0003 | 0.9302 | 0.3488 | | | |
| EG.5 | <0.0001 | <0.0001 | >0.9999 | >0.9999 | >0.9999 | | |
| EG.5.1 | <0.0001 | <0.0001 | >0.9999 | >0.9999 | >0.9999 | >0.9999 | |

| Post Delta-BA.5 RBD-heterodimer booster | | | | | | | |
|---|---|---|---|---|---|---|---|
| | Prototype | BA.4 or BA.5 | XBB.1 or XBB.1.9 | XBB.1.5 or XBB.1.9.1 | XBB.1.16 | EG.5 | EG.5.1 |
| Prototype | | | | | | | |
| BA.4 or BA.5 | >0.9999 | | | | | | |
| XBB.1 or XBB.1.9 | <0.0001 | <0.0001 | | | | | |
| XBB.1.5 or XBB.1.9.1 | <0.0001 | <0.0001 | >0.9999 | | | | |
| XBB.1.16 | 0.0080 | 0.0010 | 0.0902 | 0.5098 | | | |
| EG.5 | <0.0001 | <0.0001 | >0.9999 | >0.9999 | >0.9999 | | |
| EG.5.1 | <0.0001 | <0.0001 | >0.9999 | >0.9999 | >0.9999 | >0.9999 | |

**Figure S2. The statistical analysis of neutralizing antibody titers.**

p values were analyzed with a Kruskal-Wallis test first, and then Dunn’s multiple comparison test when the Kruskal-Wallis test was rejected.

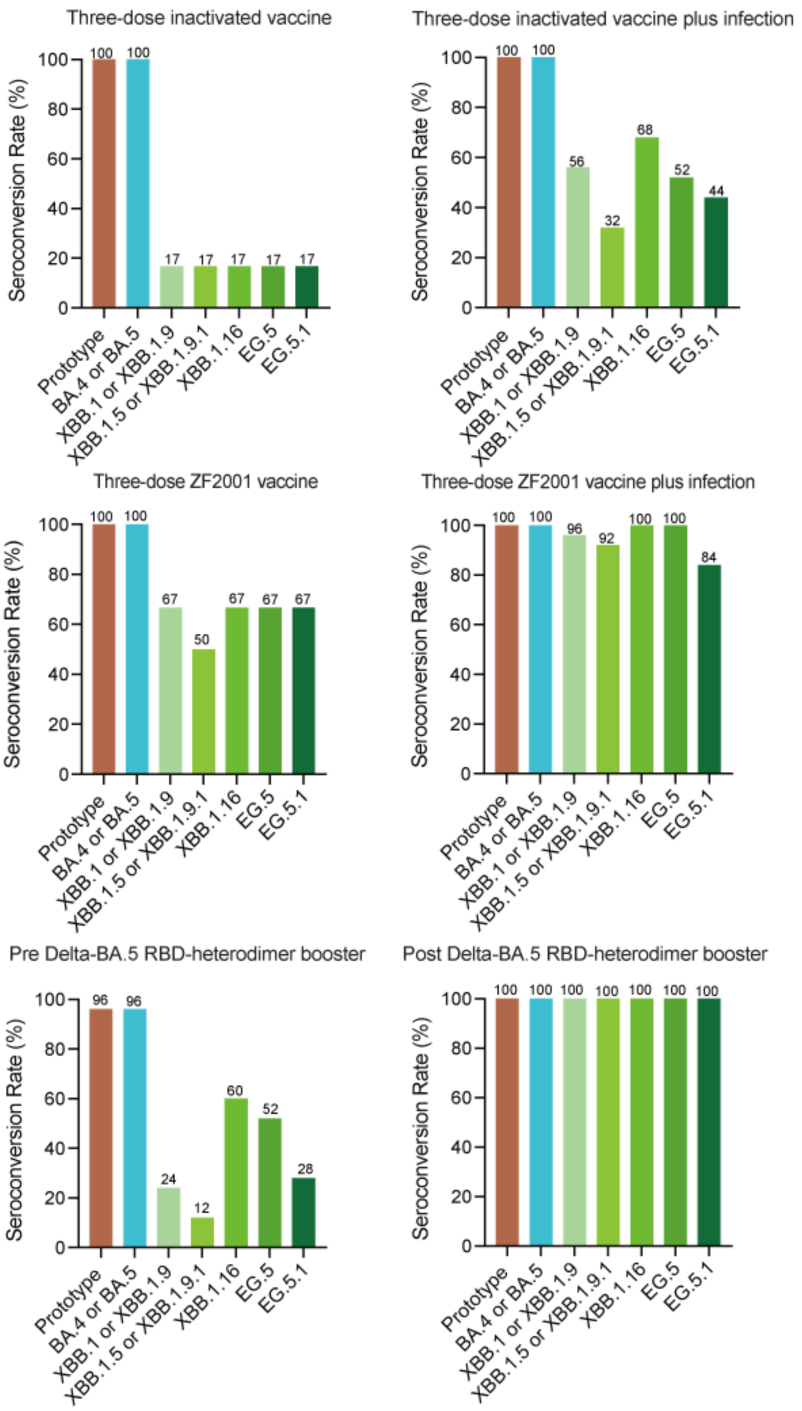

**Figure S3. Percent of neutralization positive sera against different pseudoviruses.**

The percent of neutralization positive sera was calculated as the percent of sera samples in each group with a $pVNT_{50}$ above the LOD (>1:10).

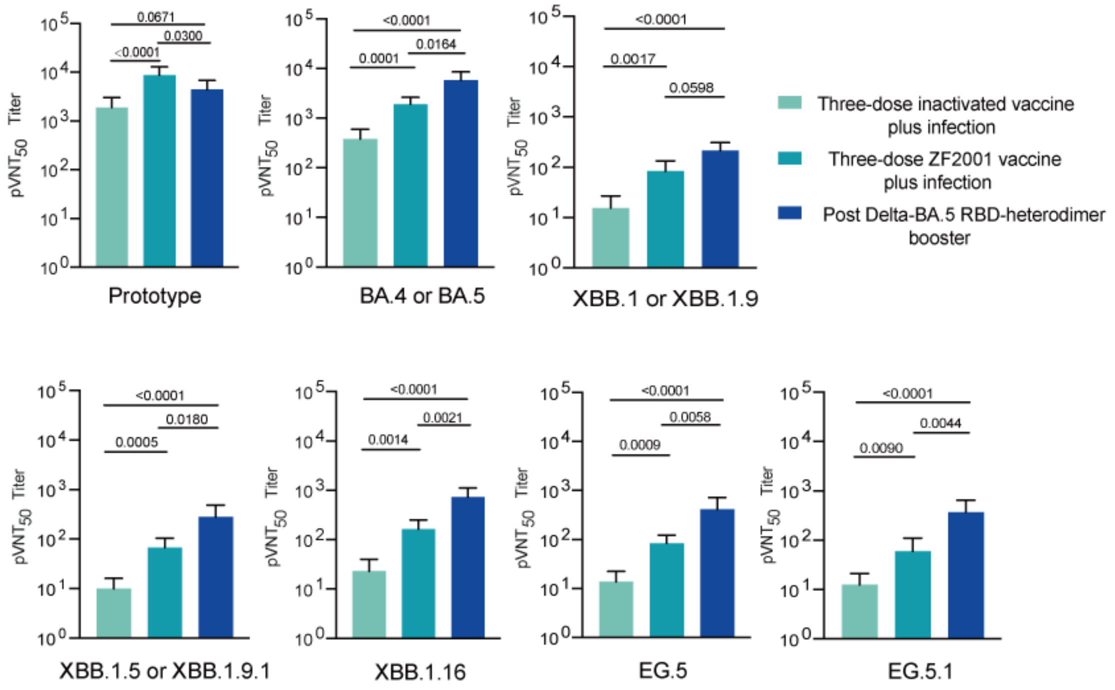

**Figure S4. The statistical analysis of neutralizing antibody titers among breakthrough infection groups and booster vaccination group.**

p values were analyzed with a Kruskal-Wallis test first, and then Dunn's multiple comparison test when the Kruskal-Wallis test was rejected.

**Acknowledgements**

This work is supported by the National Key R&D Program of China (2020YFA097100 and 2021YFC2302600) and a grant from the Bill & Melinda Gates Foundation (INV-027420). L.D. is supported by the Excellent Young Scientist Program from National Natural Science Foundation of China (NSFC) (82122031) and Youth Innovation Promotion Association CAS, China (2018113). K.X. is supported by the NSFC (82202030) and Young Elite Scientists Sponsorship Program by CAST (2022QNRC001). We thank Yuxuan Han (University of Chinese Academy of Sciences) for her assistance in the experiments.

**Author Contributions**

L.D., K.X., and G.F.G. conceived and coordinated the study. Y.A., X.Z., C.Y., and D.L. conducted the experiments. H.X. and H.H. acquired serum samples for the clinical trial. L.T. and R.W. organized the investigational products. Y.A., X.Z., K.L., L.D., and K.X. analyzed the data. K.X. and Y.A. drafted the manuscript. G.F.G. and L.D. revised the manuscript.

**Competing Interests**

Y.A., L.D, K.X., and G.F.G. are listed in the patent as the inventors of the prototype RBD-dimer as coronavirus vaccines. L.D., K.X., and G.F.G are listed in the patent as the inventors of Delta-Omicron RBD-dimer as coronavirus vaccine. L.T. and R.W. are employees at Anhui Zhifei Longcom Biopharmaceutical Co. Ltd. All other authors declare no competing interests.
